# A universal sequencing read interpreter

**DOI:** 10.1101/2022.04.16.488535

**Authors:** Yusuke Kijima, Daniel Evans-Yamamoto, Hiromi Toyoshima, Nozomu Yachie

## Abstract

Massively parallel DNA sequencing has led to the rapid growth of highly multiplexed experiments in biology. Such experiments produce unique sequencing results that require specific analysis pipelines to decode highly structured reads. However, no versatile framework that interprets sequencing reads for downstream biological analysis has been developed. Here we report INTERSTELLAR (interpretation, scalable transformation, and emulation of large-scale sequencing reads) that extracts data values encoded in theoretically any type of sequencing read and translates them into sequencing reads of any structure of choice. INTERSTELLAR first identifies sequence segments encoded in reads according to the user’s definition. Followed by error correction, values are extracted and compressed into an optimal space, which can be efficiently translated into sequencing reads of another structure. We demonstrated that INTERSTELLAR successfully extracted information from a range of sequencing reads and translated those of single-cell (sc)RNA-seq, scATAC-seq, and spatial transcriptomics to be analyzed by different software tools that have been developed for conceptually the same types of experiments. INTERSTELLAR will greatly facilitate the development of new sequencing-based experiments and sharing of data analysis pipelines.

---

In the last couple of decades, harnessing microarray and high-throughput DNA sequencing, the concept of DNA barcodes has enabled a range of pooled biological screens. Earlier examples include the establishment of the yeast deletion collection, where each strain was constructed to have two unique DNA barcodes at a deletion locus^1^. The barcoded yeast strains can be pooled and subjected to a single growth competition assay whose individual relative growth changes can be read out by barcode quantities measured by microarray or high-throughput sequencing before and after the competition^2^. This strategy has pioneered the field of chemical genomics to screen target genes of chemical compounds^3, 4^. Soon after, the same concept had also been applied to mammalian cell culture-based genome-wide gene knockdown^5^ and knockout assays^6, 7^. In such assays, cells are transduced by a lentiviral library encoding short-hairpin (sh)RNAs or CRISPR–Cas9 guide (g)RNAs. Cell growths conferred by different perturbations can be massively quantified by PCR amplification and sequencing of the small shRNA-or gRNA-encoding DNA fragments. Furthermore, experimental systems that produce chimeric fusions of distal genomic regions and those of DNA barcodes associated with different factors have enabled the exploration of chromatin conformations^8^, protein interactions^9-12^, genetic interactions^13^, and spatial cellular distribution of single-molecule RNAs^14^ in large scale. In single-cell and spatial genomics, single-cell identifiers (IDs), spatial IDs, and unique molecular identifiers (UMIs) are used to uniquely tag corresponding transcriptomes or genomic DNA fragments, which led to the development of scRNA-seq^15-18^, scATAC-seq^19, 20^, spatial transcriptomics^21, 22^, and spatial genomics^23^ technologies. The above-mentioned methods each enable multiplexing of a number of experiments at once and produce a sequencing library as a result. Sequencing libraries from different assays can also be further multiplexed for a single sequencing run by fusing an additional library-specific, unique DNA barcode(s) to each sequencing library DNA. As such, the output DNA molecules of these experiments have a range of complexities, some of which encode multiple information segments whose combinations are sometimes designed to be read out by multiple reads (e.g., paired-end reads and index reads).

However, there have been common issues—most of these sequencing-based experiments have been developed with their proprietary software tools for specific sequence read structures. While many of such tools have advanced downstream data analysis capabilities, they often cannot be reused for sequencing reads produced by other experimental systems that are developed, even for conceptually the same type of output datasets. There have been multiple “reinventions of the wheel,” requiring coding costs and labor. As seen in scRNA-seq methods, even when two similar methods have been developed with their data analysis tools, these experimental methods and tools cannot be cross-validated by exchanging them with one another. Several efforts have been made to develop flexible software tools that are capable of analyzing different read structures of a certain category of experiments, such as UMI-tools^24^, zUMIs^25^, scumi^26^ (for UMI-based RNA-seq and scRNA-seq), and SnapATAC^27^ (for scATAC-seq), but they are not effective for the ongoing development of new experiments that produce unique read structures.

Here we propose two solutions for the community: (1) the independent development of sequencing read interpreters and data analysis tools—if a read interpreter only extracts data values encoded in sequencing reads, its data analysis pipelines should be applicable for sequencing reads of other experiments that produce the same data structures; and (2) the development of a read translator—if sequencing reads of a certain format could be translated into another read structure, the existing data analysis pipelines developed for the specific read structure could be used to analyze other read structures. In this study, we have identified that these two ideas can be achieved by a single universal tool, which we have developed and called INTERSTELLAR.

## Overview of INTERSTELLAR

INTERSTELLAR interprets data values in high-throughput sequencing reads and translates them into sequencing reads of another read structure (Fig. 1). The flexible value segment identification of “source reads” enables economic development of their downstream data analysis pipelines. The translation into user-defined “destination reads” enables current data analysis tools that originally only accept a specific read structure to analyze sequencing reads of another structure.

**Figure 1.**
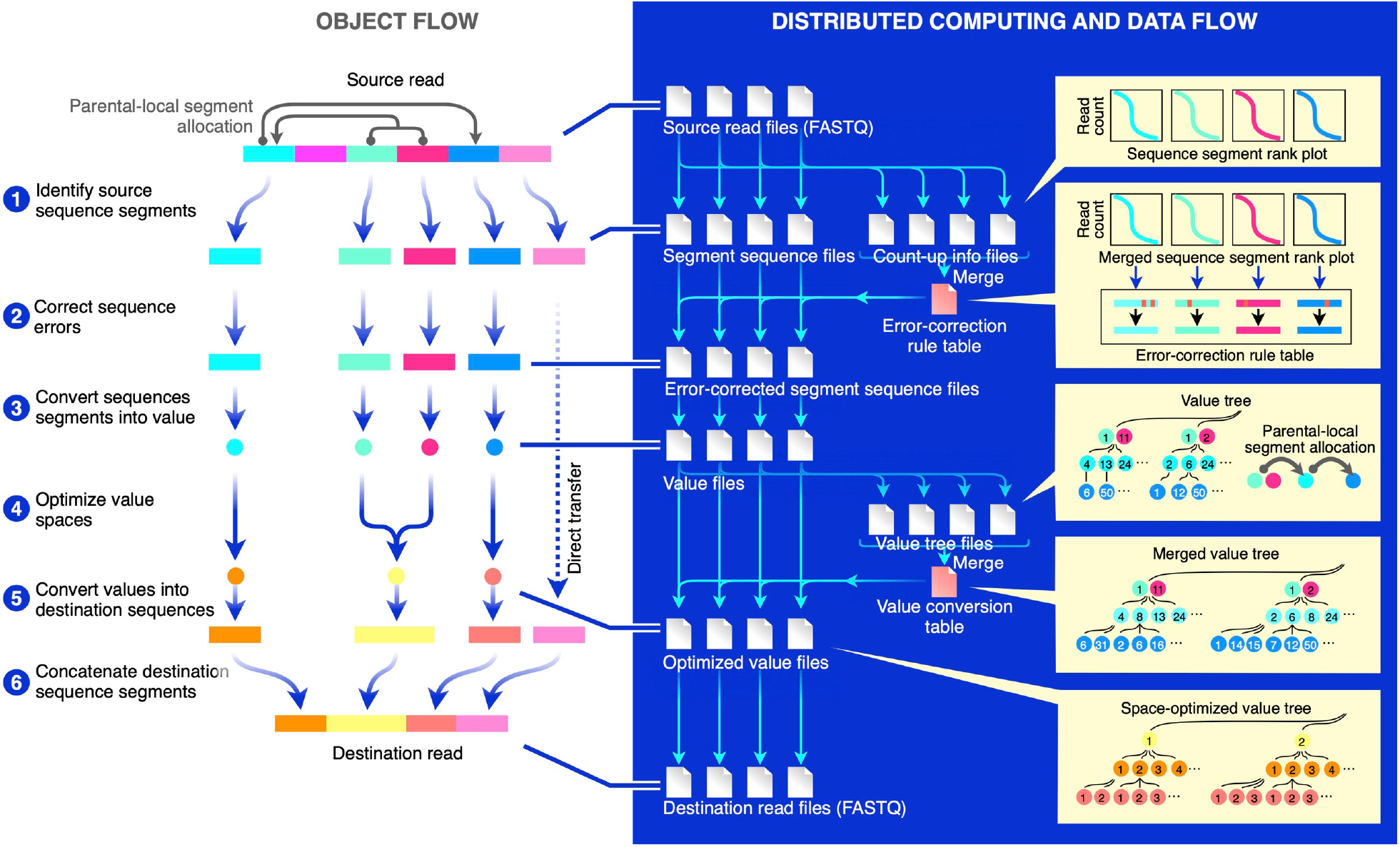
Overview of INTERSTELLAR.

INTERSTELLAR first identifies sequence segments of reads in FASTQ files according to the user’s definition provided in a process configuration file (Step 1). The source read structure can be defined flexibly, where multiple value segments on each sequencing read are specified by combinations of their lengths, locations, neighboring sequence motifs using regular expression and/or predefined allowlists, followed by identification of valid sequence segments according to their average sequencing quality (Q) scores. Three types of attributes: “combinatorial,” “parental,” and “local,” can be given to value segments with their association with other segments. A combinatorial segment group can be defined to collectively denote a specific value. A parental segment (or combinatorial parental segment group) can be paired with an independent set of local segments (or combinatorial local segment groups), where sequence-to-value interpretations of local segments are independently defined for each parental segment. For example, cell IDs and UMIs of typical scRNA-seq reads can be defined as parental segments and their local segments, respectively, where the same UMI sequences associated with different cell IDs are interpreted as different objects. Multiple source read structures can also be defined for a single set of input sequencing reads that are produced by a one-shot sequencing of different libraries.

The segment identification process can be performed independently for fragmented FASTQ files using distributed computing, where each fragmented process yields segmented sequences and count-up information for each unique segment sequence. The sequence count-up information derived from different fragmented processes is then merged to compute an error-corrected sequence for each unique segment sequence (Step 2). INTERSTELLAR enables four error correction options: “imputation-to-majority,” “mapping-to-allowlist,” Bartender^28^, and a user-developed plugin. In the imputation-to-majority correction, a merged rank-read count curve of each sequence segment is first obtained, and its knee point (the maximum curvature point) is determined. Segment sequences below the knee point are then corrected to their closest similar sequences above the knee point using the Levenshtein distance metric. Similarly, the allowlist mapping uses the Levenshtein distance metric to map input segment sequences to a user-provided allowlist. In these two options, the minor segment sequences (above the Levenshtein distance threshold) are ignored, similar to ones below Q score thresholds. The barcode sequence correction pipeline Bartender can also be employed, where input segment sequences are first grouped into clusters based on the Hamming distance metric, and minor sequences in each cluster are imputed into the top majority sequence. In contrast to the imputation-to-majority strategy, Bartender can potentially rescue valid sequences that are poorly represented in the pool. Alternatively, users can provide a shell script as a user-defined plugin to employ a customized error correction method. Once an error-correction rule table is generated, it is used to error-correct segment sequences originating from each of the fragmented FASTQ files using distributed computing. The above-mentioned read interpretation process can be applied to any high-throughput sequencing read analysis, and the generated error-corrected segment sequence files enable efficient development of their downstream data analysis pipelines.

If defined in the process configuration file, the read translation into destination read structures are next processed for the error-corrected source segment sequences (Step 3). Destination read structures can flexibly be specified by using IUPAC codes and/or allowlists of destination segment sequences. First, using distributed computing, a segment value file and a value tree are extracted from each of the error-corrected segment sequence files, where each unique segment sequence is converted into a numerical value, and parental-local segment allocations of unique values and unique combinatorial value groups are represented in a tree structure. The value tree files originating from the fragmented FASTQ files are then passed to a single computing node to generate a merged value tree. Next, the values in the merged value tree are replaced by new values to minimize the number of numerical value species for each variable in a way that they still uniquely maintain the same tree topology (Step 4). Obtaining the value conversion rule table that achieved the optimization of the merged value tree, segment value files are separately processed to derive optimized segment value files and then destination FASTQ files by distributed computing, where unique value-to-destination-sequence conversion rules are autonomously generated for the destination read structures (Steps 5 and 6). The value space optimization, which takes into account the parental-local segment allocations, is particularly effective when sequence complexities of destination segments are lower (e.g., shorter in length) than those of the corresponding source segments. This process enables a destination value sequence segment of less information representativity (vs. its source sequence segment) to host all or the maximum possible number of corresponding values represented in the source reads. (When the number of optimized values is over the information representativity of the destination value segment, frequent values are prioritized, and read information associated with any values that are not assigned to a destination segment sequence is ignored.) Throughout the process, the average Q scores of source segment sequences are bequeathed from the fragmented FASTQ files through the intermediate segment sequence and value files and given to all letters of corresponding destination segments in the generating FASTQ files. New bases that are not associated with the values inherited from the source reads are all given a Q score of 40 in the destination reads. As seen above, the distributed computing process is designed to perform many small conversion tasks in parallel, where the generation of conversion rules that require monitoring of the entire segment sequence or value space are operated using a single computing node by compressing information from each fragmented task into a small hash table data structure.

### Interpretation of highly structured barcode reads

To demonstrate that INTERSTELLAR can be used to analyze highly structured sequencing reads, we first decoded an RCP (Row-Column-Plate)-PCR library generated for massively parallel identification of BFG-Y2H^9^ barcodes. In general, RCP-PCR is developed to identify clonal DNA samples sandwiched by common PCR primer sites that are arrayed into many PCR microwell plates (Fig. 2a). Samples in each microwell plate are first amplified by forward and reverse primers with overhang sequences encoding corresponding plate row-specific barcodes (RBCs) and column-specific barcodes (CBCs), respectively. PCR products are then pooled by plates and subjected to the second round of PCR by primers with overhang sequences encoding sample plate-specific barcodes (PBCs) and Illumina sequencing adapters.

**Figure 2.**
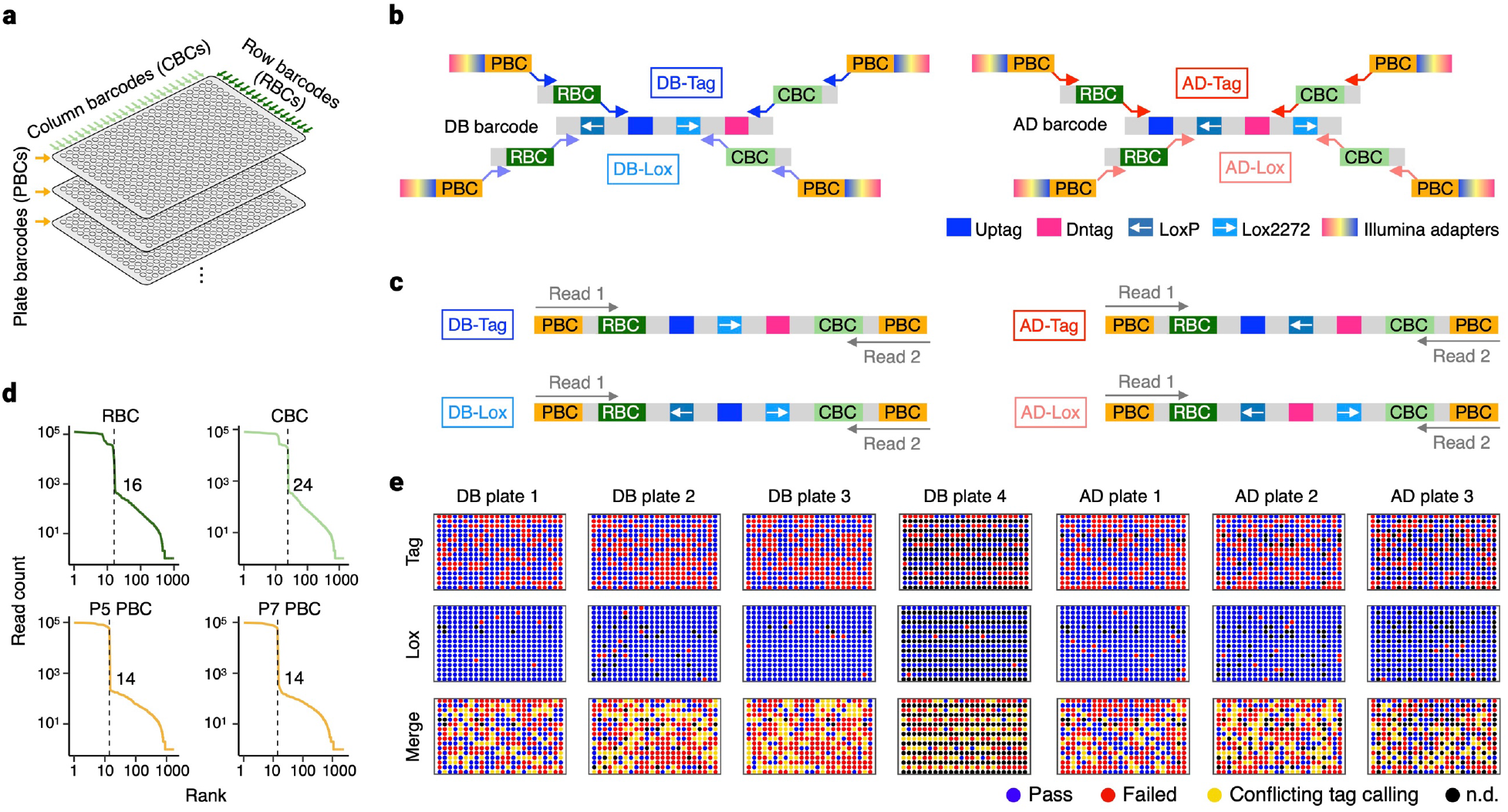
RCP-PCR analysis of BFG-Y2H barcode libraries. **a**, The conceptual diagram of RCP-PCR. **b**, Two-step PCR amplification to prepare DB-Tag, DB-Lox, AD-Tag and AD-Lox RCP-PCR libraries. **c**, Paired-end sequencing of the four library types. **d**, Rank-read count plots of RBCs, CBCs and PBCs. **e**, Identification of high-quality clonal BFG-Y2H barcode cassette samples.

BFG-Y2H is a pooled combinatorial protein interaction screening technology based on yeast two-hybrid. In this technology, DB-X MAT**a** and AD-Y MAT**α** haploid yeast strains are prepared such that each strain harbors a bait protein X fused to the DNA binding domain (DB) or a prey protein Y fused to the transcription activation domain (AD), with two X-or Y-specific barcodes and site-specific Cre recombination. The DB-X and AD-Y strains are then pooled for mating to obtain a diploid cell pool sufficiently covering the whole X-Y pair space. The diploid cell pool is then subjected to screening for cells expressing a selective marker gene due to the reconstitution of the transcription factor via interaction between X and Y. Finally, the X- and Y-specific barcodes are fused through Cre-mediated DNA recombination within cells, and the fused-barcode tags for the interacting protein pairs are identified by high-throughput sequencing.

The DB-X and AD-Y barcode cassette structures are 5’-loxP’-U1-Uptag-U2-lox2272-D1-Dntag-D2-3’ and 5’-U1-Uptag-U2-loxP’-D1-Dntag-D2-lox2272-3’, respectively, where Uptag and Dntag are barcodes to be assigned to a specific protein X or Y, U1, U2, D1, and D2 are common PCR amplification handles specific to DB-X barcodes or AD-Y barcodes, and loxP’ (reverse complement of loxP) and lox2272 are Cre recombination sites (Fig. 2b). One of the previously established methods to prepare barcoded DB-X and AD-Y strains requires subcloning of DB-X and AD-Y barcode cassettes from a pool of those with degenerated Uptag and Dntag sequences, followed by their high-throughput sequencing-based identification and quality verification employing RCP-PCR. Furthermore, to identify the clonal barcodes with high-quality base calling with a short paired-end sequencing, two different RCP-PCRs are performed against the same samples. In brief, DB-Tag RCP-PCR and DB-Lox RCP-PCR respectively identify 5’-U1-Uptag-U2-lox2272-D1-Dntag-D2-3’ and 5’-loxP’-U1-Uptag-U2-lox2272-D1-3’ regions of the same sample wells, and AD-Tag RCP-PCR and AD-Lox RCP-PCR respectively identify 5’-U1-Uptag-U2-loxP’-D1-Dntag-D2-3’ and 5’-U2-loxP’-D1-Dntag-D2-lox2272-3’ regions of the same sample wells (Fig. 2c).

To demonstrate INTERSTELLAR, DB-X barcode cassettes were subcloned into three and a half 384-well microwell plates (1,344 samples), and AD-Y barcode cassettes were subcloned into three 384-well microwell plates (1,152 samples). The identification of these barcode cassette samples required a total of 14 plate PCR reactions (4 DB-Tag, 4 DB-Lox, 3 AD-Tag, and 3 AD-Lox). The four types of RCP-PCR libraries were mixed one molar volume each and sequenced. INTERSTELLAR was used to interpret the sequencing reads of the different structures with different barcodes all at once. We first confirmed that the expected numbers of unique RBCs (16 rows), CBCs (24 columns), and PBCs (14 plates) used in this experiment were observed (Fig. 2d). Next, the sequence segment information obtained by INTERSTELLAR was analyzed by another script to identify dominantly representing (or clonal) Uptag and Dntag sequences of each well using DB-Tag and AD-Tag reads and to separately interrogate any mutational damages in loxP and lox2272 sequences of each well sample using DB-Lox and AD-Lox reads (Fig. 2e). Finally, sample wells with no high-confident agreement of Uptag sequences between DB-Tag and DB-Lox reads or Dntag sequences between AD-Tag and AD-Lox reads were discarded, yielding a total of 287 (18.7%) and 299 (26.0%) high-confident clonal DB-X and AD-Y barcode cassette samples, respectively, whose gain rates were within the range of those expected from the previous study. We also randomly selected 24 independent samples and confirmed by Sanger sequencing that 23 of their sequences were consistent with those identified by RCP-PCR.

### Translation of scATAC-seq reads

We next tested a simple translation of high-throughput sequencing reads that did not have a parental-local segment allocation. Using INTERSTELLAR, we emulated a sci-ATAC-seq dataset of Drosophila embryogenesis^29^ in 10X Genomics’ Cell Ranger ATAC, originally developed to analyze 10X scATAC-seq libraries. In sci-ATAC-seq, followed by fixation, sample nuclei are split into subpools, where open chromatin regions of nuclei are fragmented by Tn5 transposase with subpool-specific barcodes, yielding barcoded DNA fragments. After combining the barcoded nuclei samples, they are split into subpools again, where unique combinations of Illumina i5 and i7 indexed adapters are concatenated to the fragmented DNA in the nuclei of the corresponding subpool by PCR. Finally, PCR products are pooled for a single Illumina sequencing run. This multistep split-and-pool strategy is designed to tag open chromatin DNA fragments of massive single cells with cell-specific combinations of DNA barcodes. Each library was sequenced by a total of four reads: two paired reads to sequence the genomic region and two index reads each to identify a combination of four barcodes (Fig. 3a). On the other hand, 10X scATAC-seq is an emulsion-based method that encapsulates single cells in water-in-oil droplets with unique droplet-specific 16-bp barcodes. In each droplet, open chromatin regions are fragmented by Tn5 and concatenated to the droplet-specific barcodes (i.e., cell IDs), each of whose products is sequenced by a total of three reads (Fig. 3a).

**Figure 3.**
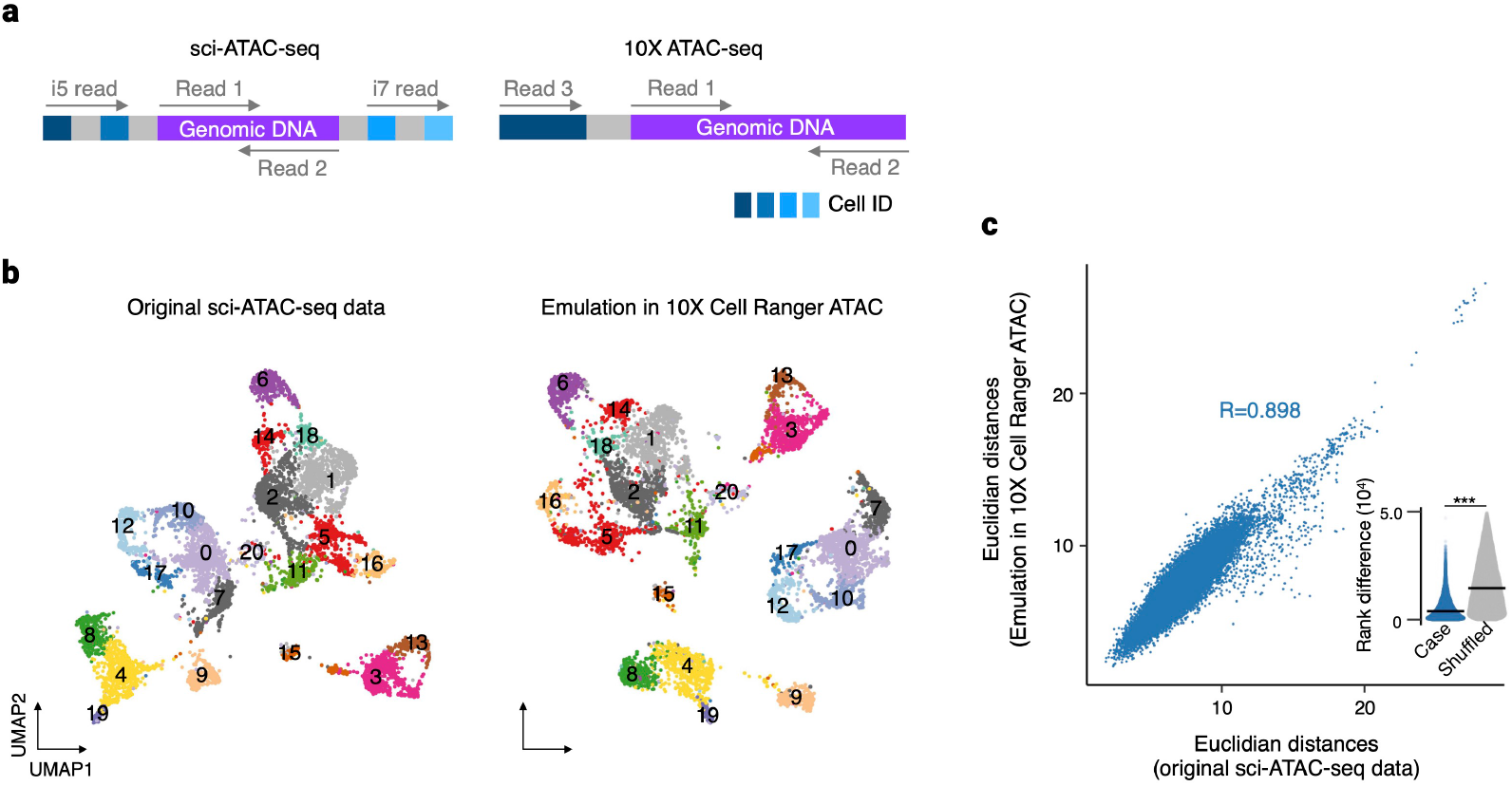
Translation of sci-ATAC-seq reads for 10X Cell Ranger ATAC. **a**, Read structures of sci-ATAC-seq and 10X scATAC-seq. **b**, Two-dimensional UMAP embeddings of sci-ATAC-seq data processed by its original pipeline for Drosophila embryo 6 to 8 hours after egg laying, and that obtained by Cell Ranger ATAC with the read translation using INTERSTELLAR. Cell state annotations obtained by the original pipeline were applied to both embeddings. **c**, Correlation in distance of two cells between the high-dimensional genomic accessibility count space of the original sci-ATAC-seq data and that by Cell Ranger ATAC. For each dataset, Euclidean distances in a high-dimensional LSI space were measured for the same 50,000 randomly sampled cell pairs. The inset sina plot represents rank difference distribution in the Euclidean distance of the same cell pairs before and after translation. The crossbar represents the median. \*\*\**P*-value < 2.2e–16.

We translated the sci-ATAC-seq dataset of embryos 6 to 8 hours after egg laying to the read structure of 10X scATAC-seq, performed two-dimensional uniform manifold approximation and projection (UMAP) embeddings of the high-dimensional single-cell genomic accessibility count matrix by Cell Ranger ATAC, and compared it with that from the original sci-ATAC-seq reads obtained using the proprietary data analysis pipeline (Fig. 3b). The cell state clusters identified by the original sci-ATAC-seq pipeline were markedly replicated in the translated dataset analyzed by Cell Ranger ATAC. To assess the data similarity between the original and emulated datasets, we compared Euclidean distances in a high-dimensional latent semantic indexing (LSI) space between randomly selected pairs of cells in the two datasets and found a Pearson’s correlation coefficient of 0.898 (Fig. 3c). Furthermore, we also measured the rank difference in the Euclidean distance of the same cell pairs in the two datasets and compared it with the random expectation (see Methods). We demonstrated that the data profiles were significantly preserved after the read translation (*P*-value < 2.2e–16). Furthermore, we examined if the read translation by INTERSTELLAR maintained the ability of the dataset to have its biological information extracted, similar to the original pipeline. The sci-ATAC-seq reads of three embryonic samples of 2 to 4, 6 to 8, and 10 to 12 hours after egg laying were pooled, translated, and analyzed by Cell Ranger ATAC. We confirmed that the analysis successfully recaptured the dynamic diversification of single-cell genomic accessibilities through Drosophila embryogenesis and cell state-specific marker gene accessibilities and their genomic distributions (Extended Data Fig. 1).

### Cross-evaluation of different scRNA-seq reads and software tools

Differences in information capacity (or sequence representativity) between source segments and destination segments need to be taken into consideration in some read translations. For example, when the total base-pair length of a destination segment is shorter than that of the corresponding source segment(s), the destination segment might not be able to represent all the values observed in the source reads. However, the value space optimization implemented in INTERSTELLAR greatly alleviates this issue by allowing the end-user to interpret parental-local segment allocations of the source read structure. For example, UMIs of typical scRNA-seq reads are local to their corresponding transcription products of corresponding single cells that are uniquely encoded in other segments. Some scRNA-seq libraries employ shorter UMI segments than others, but usually, even the read translations from the latter to the former do not have major issues, because the number of unique UMI sequences observed for each of the combinatorial parental segments is practically limited and the value space for the UMIs representing the entire scRNA-seq data can be largely compressed.

To demonstrate that different scRNA-seq read structures can be practically translated to and from each other, as well as analyzed by different software tools of choice that have originally been developed for specific read structures, we obtained four sequencing read datasets of 10X Chromium V3^30^ (mouse heart; 16-bp cell barcode and 12-bp UMI), Drop-seq^15^ (mouse eye; 12-bp cell barcode and 8-bp UMI), Quartz-seq2^31^ (mouse stromal vascular fraction; 14-bp cell barcode and 8-bp UMI), and SPLiT-seq^17^ (mouse brain; three combinatorial 8-bp cell barcodes and 10-bp UMI), all of which are representative state-of-the-art scRNA-seq methods (Fig. 4a). We compared their analysis results obtained using the original software tools with those by 10X Cell Ranger (originally developed for 10X Chromium) and dropseq-tools (originally developed for Drop-seq) with read translation using INTERSTELLAR (Fig. 4b).

**Figure 4.**
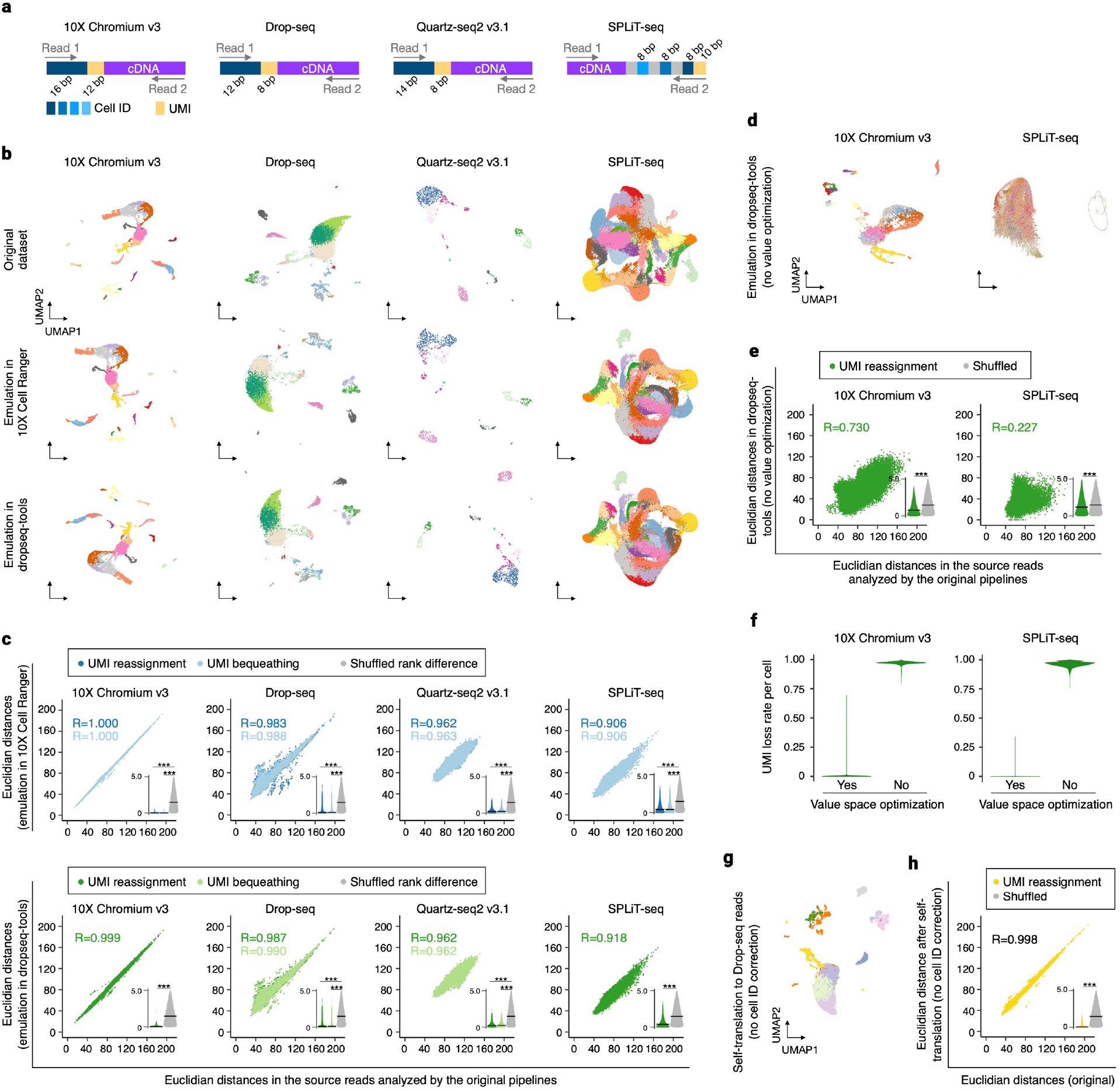
Translation of scRNA-seq reads for different analysis pipelines. **a**, Read structures of different scRNA-seq methods. **b**, Two-dimensional UMAP embeddings of scRNA-seq datasets processed by their original pipelines and those analyzed using 10X Cell Ranger and dropseq-tools by read translation using INTERSTELLAR with the “UMI reassignment” strategy. Cell state annotations obtained by the original pipelines were applied to the translated results. **c**, Correlation in distance of two cells between the high-dimensional transcriptome spaces of the original datasets and those translated for Cell Ranger and dropseq-tools with the “UMI reassignment” and “UMI bequeathing” strategies. For each dataset, Euclidean distances in the gene expression count matrix were measured for 50,000 randomly sampled cell pairs. The bottom-right inset sina plot) of each panel represents rank difference distribution in the Euclidean distance of the same cell pairs before and after translation. The crossbar represents the median. **d**, Two-dimensional UMAP embeddings of 10X Chromium and SPLiT-seq datasets processed by their original pipelines, and those analyzed by dropseq-tools using INTERSTELLAR without value space optimizations. **e**, Correlation in distance of two cells between the high-dimensional transcriptome spaces of the original datasets and those translated for dropseq-tools without value space optimizations. **f**, UMI loss rate per cell with and without value space optimizations. **g**, Two-dimensional UMAP embedding of the Drop-seq dataset self-translated for dropseq-tools with no cell ID error correction. **h**, Correlation in distance of two cells between the high-dimensional transcriptome spaces of the original and self-translated Drop-seq datasets. \*\*\**P*-value < 2.2e–16.

While all of the read translations were first performed with the value space optimization where UMI sequences locally to the cell ID segments were optimized and translated to destination segment sequences, we also examined another emulation strategy that bequeathed the same UMI sequences to the destination reads for the translations whose destination UMI lengths were equal to or longer than those of the sources (constant sequences were added to the source UMIs to meet the length of the destination UMIs). In these sequencing library emulations, 10X Chromium V3 and Drop-seq read datasets were also self-translated by INTERSTELLAR for 10X Cell Ranger and dropseq-tools, respectively. The transcriptomic profiles of single cells in the translated datasets were compared to those in the original datasets by their correlations in the Euclidean distances of two cells in the high-dimensional transcriptome space and by the rank difference distribution of the Euclidean distances in the two datasets compared to the random expectation (Fig. 4c). The “UMI reassignment” (value space optimization) and “UMI bequeathing” strategies both conferred almost identical results. Furthermore, the read translation that required shortening of the UMI lengths with the UMI reassignment strategy also demonstrated similar results to those which did not require UMI shortening, suggesting efficient read translations. Indeed, 98.17% and 99.99 % of single cells retained the complete source UMI segment values in the emulation of 10X Chromium and SPLiT-seq datasets for dropseq-tools, respectively, whereas the translations without value space optimizations showed markedly poor cell state preservations (Fig. 4d) and Euclidean distance correlations (Fig. 4e), and no single cells retained complete UMI segment values (Fig. 4f).

While all of the read dataset translations by INTERSTELLAR largely retained the single-cell transcriptome profiles of the original datasets processed by their proprietary tools, the self-emulation of the Drop-seq dataset for dropseq-tools showed a variance in contrast to that of the 10X Chromium V3 for 10X Cell Ranger. This was likely due to the use of the Levenshtein distance-based error correction for interpretation of the cell ID segments in INTERSTELLAR, where 10X Cell Ranger also employs a Levenshtein distance-based error correction, but dropseq-tools (Drop-seq) employs a unique error correction metric trained by its cell ID synthesis errors. (Quartz-seq pipeline (Quartz-seq2) and splitseq-tools (SPLiT-seq) employ Sequence-Levenshtein distance^32^ and Hamming distance, respectively.) To test this hypothesis, we performed self-emulation of the Drop-seq library by the UMI reassignment strategy with no error correction in the cell ID segments (Fig. 4g) and demonstrated that a much smaller variance was produced when the error correction process was fully performed in dropseq-tools (Fig. 4h). Apart from the variance observed in the self-translation, generally, the other sources of variance between the emulated and original data could be explained by differences in other software-specific downstream data processes that are independent of INTERSTELLAR. For example, the Quartz-seq pipeline only obtains read count profiles of coding regions, while Cell Ranger and dropseq-tools account for 3’ UTR regions.

To further challenge INTERSTELLAR with translating a more complex read structure with high order value space optimizations, we translated a read pool of 10X Chromium multiplexed into a hypothetical destination read structure with multi-layered parental-local segment allocations and translated them back to the 10X Genomics Chromium read structure (Extended Data Fig. 2). While the multiplexed 10X Chromium libraries are sequenced by a combination of two reads, one for cell ID and UMI and the other for transcripts, and another combination of two reads for sample indices, we designed the hypothetical destination read structure to have a total of 17 segment variables in addition to the transcript sequence segment. Analyzing both the original and “round-tripped” reads using 10X Cell Ranger, we confirmed that INTERSTELLAR could well preserve the single-cell transcriptome profile information.

### Translation of spatial transcriptomics reads

In the translation of scATAC-seq and scRNA-seq reads, the values extracted from the source segment sequences were assigned to destination segment sequences that were arbitrarily generated or selected from a given allowlist. However, in INTERSTELLAR, the user can also define the translation of source segment sequences to corresponding destination segment sequences by providing a sequence conversion table. To demonstrate the use of this function, we translated spatial transcriptome reads of Slide-seq^21^ to the read structure of 10X Visium^22^ and analyzed them by Visium’s proprietary software Space Ranger. The two spatial transcriptomics technologies have been developed with similar conceptual designs. In brief, a tissue sample section is applied to a surface material where reverse transcription (RT) primers with unique positional barcodes are immobilized on distinct locations of the two-dimensional surface. Transcriptomes leaked from cells of specific positions are captured by proximal, positionally barcoded RT primers and reverse transcribed such that they are fused to the positional barcodes for pooled high-throughput sequencing. Both read structures are similar to those of scRNA-seq (i.e., UMI employed to uniquely identify each RT primer molecule; Fig. 5a). The significant difference between the two spatial transcriptomics technologies is in the preparation of positionally barcoded RT primers. In Visium, RT primers with known positional barcode sequences are immobilized on predetermined spots of the two-dimensional surface. In contrast, Slide-seq employs uniquely barcoded beads of unidentified barcode sequences, distributes them onto a glass slide surface, and retrospectively identifies the positional barcode sequences by sequencing-by-ligation before applying sample tissue.

**Figure 5.**
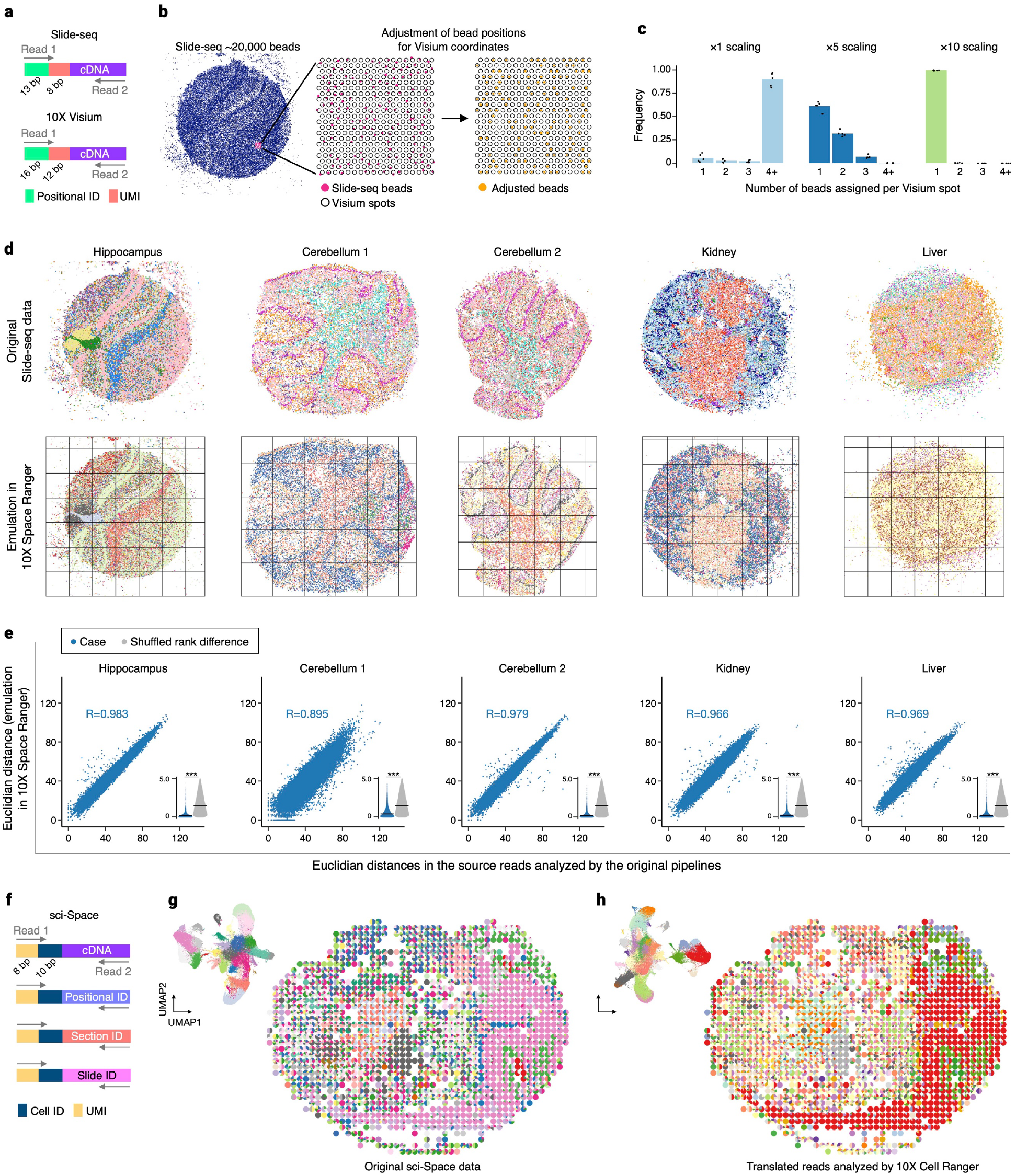
Translation of spatial transcriptomics reads. **a**, Read structures of Slide-seq and 10X Visium. **b**, Strategy to associate Slide-seq positional barcodes to those of multiple 10X Visium slides. Multiple Visium slides are first tiled across an enlarged Slide-seq field with a given scaling factor. Slide-seq positional barcodes are then associated to the closest Visium positional barcodes. **c**, Relative frequency distributions in number of Slide-seq positional barcodes assigned per Visium positional barcode with scaling factors of ×1, ×5, and ×10. **d**, Original Slide-seq datasets and those analyzed by 10X Space Ranger with ×10 scaling. Each grid represents a tiled Visium slide.

We analyzed five Slide-seq libraries (Hippocampus, Cerebellum 1, Cerebellum 2, Kidney, and Liver) using Space Ranger. While 10X Visium and Space Ranger are designed to analyze 4,992 spatial spots at a time, a Slide-seq slide is usually composed of more than 20,000 barcoded beads. Therefore, we scaled the Slide-seq sample space coordinates and tiled the Visium slides, where each tile was treated as a single Visium experiment, and Slide-seq bead positions were further aligned to the most proximal Visium spots of the corresponding tiles (Fig. 5b). We first sought the best scaling size of the Slide-seq slide that enabled the assignment of single Slide-seq beads to unique Visium spots. Among the three expansion scales of ×1, ×5, and ×10, we found that the ×10 scaling enabled an average of 99.6 % Slide-seq positional barcodes across the five samples to find their unique Visium spots (Fig. 5c). After generating a sequence conversion table for Slide-seq positional barcodes to their corresponding Visium spot barcode of corresponding tiles, reads were translated using INTERSTELLAR and analyzed by Space Ranger. As a result, the spatial gene expression patterns obtained by the read translations were markedly similar to those analyzed by the original pipeline (Fig. 5d). Euclidean distances of two positional pairs in the high-dimensional transcriptome space were highly correlated before and after the read translation (Fig. 5e). The median rank differences of Euclidean distances before and after read translation were all significantly lower than those of random expectation but seemed highly dependent on the sequencing quality. The median rank differences before and after translation for Hippocampus, Cerebellum 1, Cerebellum 2, Kidney and Liver were 1,479, 3,874, 1,626, 2,054 and 1,944, respectively, where their average read counts per positional barcode were 5,363, 315, 10,262, 9,544, and 15,481, respectively.

Recent studies have employed polyadenylated cellular RNA barcodes for scRNA-seq to obtain single-cell transcriptomic information together with cell clone barcodes (e.g., CellTagging^33^ and LARRY^34^), cell lineage barcodes (e.g., scGESTALT^35^) and genetic perturbation reagent information (e.g., Perturb-seq^36^ and CROP-seq^37^). To demonstrate that INTERSTELLAR can analyze such multimodal sequencing read datasets, we also translated the recently published sci-Space^38^ reads for 10X Cell Ranger. The sci-Space read structure is composed of Read 1, always encoding cell ID and UMI segments, and Read 2, encoding either cDNA sequence, spatially deposited positional ID, section ID, or slide ID (Fig. 5f), where a combination of the three IDs paired with the same cell ID defines a predefined two-dimensional coordinate among multiple sci-Space slides for that single cell. The cell states identified from the translated reads using Cell Ranger and their spatial positions were markedly similar to those of the original reads analyzed by the proprietary tool (Fig. 5g and h).

## Discussion

As supported by the present demonstrations, INTERSTELLAR can theoretically interpret sequencing reads of any structure and benefit any new high-throughput sequencing-based method development. We performed read translations and data analyses using different software tools for scATAC-seq, scRNA-seq, and spatial transcriptomics reads and compared the results with those from the original reads analyzed by the original proprietary software tools. Although the overall results were markedly similar between the original and emulated results, there were different levels of the variances observed. The difference in the results can be explained by three potential sources: (1) the read interpretation process, (2) the destination segment assignment process, and (3) differences in the value analysis processes between different software tools, where INTERSTELLAR is responsible for the first two. From the demonstrations of translating scRNA-seq reads, the error correction step of the read interpretation process was suggested to be a potential major source of the variance seen, in which the error correction of the read interpretation was likely to make the error correction steps implemented in different software tools ineffective (i.e., overriding of the error correction strategy by INTERSTELLAR). Although the Levenshtein distance metric is the default for the non-allowlist-based error correction of INTERSTELLAR, and this is practically not an issue for most sequencing read data analyses, it can be replaced with Bartender or a user-developed plugin. The destination segment sequence assignment process is the only potential source of the loss of information encoded in the source reads when the information capacity (or representativity) of a destination segment is less than that of the corresponding source segment. To address this issue, we implemented theoretically the best value space optimization strategy that utilizes the user-defined information of parental-local segment allocations and successfully demonstrated that the information loss could be minimal for the read translations with a reduction in sequence representativity.

In the last couple of decades, beyond the (epi)genomic and transcriptomic analyses of clinical samples and various species, applications of massively parallel short-read sequencing technologies have enabled the development of wide-ranging biological assays, and the field continues to expand rapidly. While it has been a practice to develop and combine proprietary sequencing read interpreters and data analysis pipelines with the development of new sequencing-based assays, we propose a shift to a new form, where the community uses a common sequencing read interpretation and translation platform, like INTERSTELLAR, develops only the data analysis parts and shares them separately for the best utilization of data processing resources.

## Acknowledgments

We thank the members of the Yachie laboratory at the University of British Columbia and the University of Tokyo for useful comments and discussions throughout the course of this study, especially Samuel King for proofreading the manuscript. This study was performed under the Canada Research Chair program supported by the Canadian Institutes for Health Research (CIHR) and the Pilot Innovation Fund (PIF003) supported by Genome British Columbia to N.Y. Y.K. and D.E-Y. were supported by the JSPS DC2 and DC1 Fellowships, respectively, from the Japan Society for the Promotion of Science (JSPS). Sequencing read analysis was performed using the SHIROKANE Supercomputer at the University of Tokyo Human Genome Center.

## Author contributions

Y.K. and N.Y. conceived the study and designed INTERSTELLAR. Y.K. implemented INTERSTELLAR. D.E-Y performed the RCP-PCR experiments. Y.K. performed the data analyses. H.T. supported the data analyses. Y.K. and N.Y. wrote the manuscript.

## Methods

### Datasets

scATAC-seq, scRNA-seq, and spatial transcriptomics datasets obtained for this study are listed in Supplementary Table 1. For the sci-ATAC-seq dataset, we found only FASTQ files where cell IDs are recorded in the header line of each entry. Therefore, we generated new FASTQ files such that each sequencing read entry encoded the cell ID with per base Q scores of 30. Similarly, for the Slide-seq datasets, we extracted positional barcodes, UMIs, and cDNA sequences from the available BAM files and regenerated FASTQ files by setting the per base Q scores of positional barcodes and UMIs to 37. Q scores for the cDNA sequences were inherited from the BAM files.

### Preparation of barcoded plasmids for RCP-PCR

DB-X and AD-Y barcode plasmids were respectively constructed from pDN0510 and pDN0509^39^ by assembling BFG-Y2H DB-X and AD-Y barcode fragment pools^9^ by three-fragment Gibson DNA assembly^40^ as follows: DB-X Uptag and Dntag fragment pools were amplified using the random barcode templates DB-BC-UP and DB-BC-DN, with the primer pairs DB-BC-UP_F/DB-BC-UP_R and DB-BC-DN_F/DB-BC-DN_R, respectively. Similarly, AD-Y Uptag and Dntag fragment pools were amplified using the random barcode templates AD-BC-UP and AD-BC-DN, with the primer pairs AD-BC-UP_F/AD-BC-UP_R and AD-BC-DN_F/AD-BC-DN_R, respectively. Each PCR was performed in a 35 µl volume, including 3.6 µl of 10 pM template, 0.7 µl each of 10 µM primers, 0.2 µl Phusion DNA Polymerase, 7 µl 5× Phusion HF detergent-free Buffer (Thermo Fisher Scientific, F520L), and 0.28 µl of 25 mM dNTPs with the following thermal cycler conditions: 98 °C for 30 s, 5 cycles of 98 °C for 10 s, 65 °C for 10 s, and 72 °C for 10 s, and then 24 cycles of 98 °C for 10 s and 72 °C for 10 s, followed by 72 °C for 5 mins for the final extension. The pDN0509 and pDN0510 backbones were linearized by PI-PspI (NEB) following the manufacturer’s instruction. Each Gibson DNA assembly of the Uptag pool, Dntag pool and linearized backbone was performed in a total of 20 µl volume with 25 fmol of each of the backbone and barcode fragments. The reaction was incubated at 50 °C for 1 hour, and 1 µl was used to transform 50 µl of One Shot(tm) ccdB Survival(tm) 2 T1R Competent Cells (Thermo Fisher Scientific, A16460) according to the manufacturer’s instructions. The transformation samples were spread on 245 mm × 245 mm square LB+ampicillin plates and incubated overnight at 37 °C for colony isolation. Single colonies were picked by QPix 450 robot (Molecular Device) and arrayed into 384-well plates with liquid LB+ampicillin media. Oligonucleotides used in this protocol are listed in Supplementary Table 2.

### RCP-PCR

RCP-PCRs were performed as described previously^9^. In brief, the first Row-Column PCRs (RC-PCRs) for DB-Tag, DB-Lox, AD-Tag, and AD-Lox fragments were performed with the primer sets encoding Row or Column identifiers (Supplementary Table 2). Each RC-PCR was performed in 10 µl volume, including 4 µl of 16-fold diluted overnight culture as templates, 1 µl each of 2 µM primers, 0.2 µl Phusion DNA Polymerase, 2 µl 5× Phusion HF Buffer (NEB), and 0.08 µl of 25 mM dNTPs. The thermal cycler conditions for DB- and AD-Tag RC-PCRs were: 95 °C for 3 mins, 30 cycles of 95 °C for 10 s, 63 °C for 10 s, and 72 °C for 15 s, and then 72 °C for 5 mins for the final extension. The conditions for DB- and AD-Lox RC-PCRs were: 95 °C for 3 mins, 30 cycles of 95 °C for 10 s, 66 °C for 10 s, and 72 °C for 15 s, and then 72 °C for 5 mins for the final extension. (Note that 4× 96-well PCRs were performed for each 384-well template sample plate for better sample handling.) The RC-PCR products were pooled by plates and purified using FastGene PCR purification kit (Nippon Genetics) and subjected to plate PCR (P-PCR) using custom indexed primers for Illumina library preparation listed in Supplementary Table 2. Each P-PCR was performed in 40 µl volume including 2× Phusion High Fidelity PCR Master Mix (NEB), 1 µl each of 10 µM Forward and Reverse plate primers, and 1 ng of size selected RC-PCR product with the following thermal cycler condition: 98 °C for 30 s, 15 cycles of 98 °C for 10 s, 60 °C for 10 s, and 72 °C for 1 min, and then 72 °C for 5 mins for the final extension. The P-PCR products were pooled and quantified by qPCR using the KAPA Illumina library quantification kit (Kapa Biosystems) and sequenced by MiSeq (Illumina, 2×250 bp paired-end sequencing).

### Interpretation of RCP-PCR reads

Using INTERSTELLAR, we identified Uptag, Dntag, loxP, and/or lox2272 segments of DB-Tag, DB-Lox, AD-Tag and AD-Lox reads with RBCs, CBCs and PBCs. We discarded any sequence segments whose average Q scores were below 20 or whose minimum per base Q scores were below 10. P5 PBCs, P7 PBCs, RBCs, and CBCs were error-corrected using the allowlists with a maximum Levenshtein distance threshold of 1. The process configuration file of INTERSTELLAR is available at https://github.com/yachielab/Interstellar/blob/main/config/fig2_RCPPCR/rcppcr.conf.

### Identification of high-quality clonal barcode cassettes

To determine sample wells containing clonal barcode cassettes, we analyzed the RCP-PCR data interpreted by INTERSTELLAR with the following criteria. For each well, we first determined the most dominant Uptag and Dntag in the Tag RCP-PCR result. If the occupancies of the most frequent tags were both above 50%, the Uptag and Dntag were regarded as clonal in the well. The validities of loxP and lox2272 were separately evaluated using the reads from the Lox RCP-PCR, with the criterion of 70% or more reads encoding the correct sequences. Since DB- and AD-Lox RCP-PCR reads respectively encode Uptags and Dntags, we also determined the most dominant Uptag or Dntag from the Lox RCP-PCR reads of each well with the same criterion used to call the clonal Uptags and Dntags from the Tag RCP-PCR reads. Finally, wells with single dominant Uptag and Dntag pairs, valid loxP and lox2272 sequences, and no conflict in Uptag or Dntag between Tag and Lox RCP-PCRs were called to contain high-quality clonal barcode cassettes. The Python script for this process is available at https://github.com/yachielab/Interstellar/blob/main/utils/analyze_rcppcr.py. For validation, we randomly selected 24 wells of predicted clonal barcode cassettes, cultured the corresponding bacterial samples overnight in LB+ampicillin media at 37 °C, extracted plasmids using FastGene plasmid kit (Nippon Genetics), and performed Sanger sequencing with Term-Rvs primer (Supplementary Table 2).

### Translation of sci-ATAC-seq reads

Using INTERSTELLAR, we identified four combinatorial cell IDs and a genomic DNA region of each sci-ATAC-seq read combination and translated them into the read structure of 10X scATAC-seq. We discarded any read containing genomic DNA segments whose average Q scores were below 20 or whose minimum per base Q scores were below 5. Cell IDs in the source reads were error-corrected with the maximum Levenshtein distance threshold of 1 using a position-specific cell ID allowlist. Each combination of cell IDs was translated into a 16-bp barcode selected from the cell ID allowlist of 10X Chromium scATAC-seq. The process configuration file of INTERSTELLAR is available at https://github.com/yachielab/Interstellar/blob/main/config/fig3_sciATAC/sciATAC_to_10xATAC.conf.

### scATAC-seq data analysis

We first analyzed the sci-ATAC-seq reads of Drosophila single cells for 6 to 8 hours after egg laying (GSE101581) by using 10X Cell Ranger ATAC (version 1.2.0) with read translation using INTERSTELLAR. FlyBase version R6.25 and Ensemble BDGP6.95 were used as a reference genome and for genomic annotation, respectively, to obtain a genomic accessibility count matrix. For comparison, we obtained the original raw genomic accessibility count matrix (2-kbp bins across the genome; GSE101581) produced in the original study. Following the workflow used in the original study, cells with the lowest 10% read counts were discarded, resulting in 7,092 cells. Furthermore, the genomic accessibility count matrix was limited to the top 20,000 2-kbp bins of frequently mapped reads across cells for the subsequent steps. After obtaining the genomic accessibility count matrix from the translated 10X scATAC-seq reads using Cell Ranger ATAC, the following analyses were limited to the 7,092 cells observed in both matrices. Both genomic accessibility count matrices were processed by Signac version 1.0.0^41^. For each matrix, accessibility count normalization was performed by RunTFIDF(), and the normalized matrix was processed by RunSVD() for low-dimensional data projections by Singular Value Decomposition (SVD). After identifying the top 30 LSI components, LSIs correlated with single-cell read depth with Pearson’s correlation coefficients of over 0.5 were removed (LSI components 1 and 4 and components 1 and 5 were removed from the original and translated datasets, respectively). The remaining 28 LSI components were used for UMAP embedding of the data using RunUMAP() to identify cell clusters using k-nearest neighbor (kNN) clustering by FindNeighbors() and FindClusters() with a resolution parameter of 1. We also analyzed the sci-ATAC-seq reads of Drosophila single cells for all of the available developmental stages in the same study (2 to 4, 6 to 8, and 10 to 12 hours after egg laying) by using Cell Ranger ATAC with read translation. The data analyses by Cell Ranger ATAC were first independently performed for three stage-specific samples—each sample with the same reference genome and genomic annotation. We aggregated the genomic accessibility count matrices from all samples into a single matrix using Cell Ranger ATAC and analyzed it by Signac. Low-quality cells were discarded according to the instruction of Signac; the nucleosome signal scores and transcription start site (TSS) enrichment scores of cells were computed by NucleosomeSignal() and TSSEnrichment(), respectively, and cells with nucleosome signal scores of <2, TSS enrichment score of >2, and %reads mapped to identified accessibility peaks of >40 were retained. Furthermore, identified accessibility peaks with 200–100,000 mapped reads across retained single cells were used to construct the high-quality genomic accessibility count matrix, followed by read count normalization and low-dimensional data projection as described above.

### Comparison of the original and translated high-dimensional datasets

To compare two high-dimensional count matrices obtained by applying different data processing methods to the same scATAC-seq, scRNA-seq, or spatial transcriptome read dataset, we adopted the following metric. First, 50,000 pairs of high-dimensional data points (e.g., transcriptome profiles of single cells or spatial positions) were randomly sampled, and their Euclidean distances in the two datasets were compared. Furthermore, to quantitatively evaluate the similarity of the two datasets in a nonparametric manner, we defined the rank difference *ΔR*_*i,j*_ of the same high-dimensional data pairs (*i, j*) between the two datasets as follows and evaluated their distribution compared to that obtained from two data pairs each independently sampled from the two datasets:

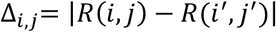

where *i* and *j* are randomly sampled high-dimensional data points in a set *θ* (*i,j* ∈ *θ*), *i’* and *j’* are the corresponding data points in a projected (translated) set *θ’* (*i’,j’* ∈ *θ’*), and *R*(*x,y*) represents the rank of Euclidean distance of *x* and *y* (*x,y* ∈ *X*).

### Translation of scRNA-seq reads

10X Chromium V3 scRNA-seq, Drop-seq, Quartz-seq2 v3.1, and SPLiT-seq reads were analyzed by INTERSTELLAR to identify their cell ID(s), UMI, and cDNA sequence segments. We discarded sequencing reads with any segment whose average Q score was below 20 or whose minimum per base Q score was below 5. The cell IDs of Drop-seq reads were corrected by the imputation-to-majority correction with the maximum Levenshtein distance threshold of 1. For 10X Chromium V3, Quartz-seq2 and SPLiT-seq reads, the cell IDs were corrected using allowlists with the maximum Levenshtein distance threshold of 1. In the read translation for Cell Ranger, the cell ID values and UMI values were assigned to sequence segments selected from the whitelist of 10X Chromium V3 and 12-bp random sequence segments, respectively. In the read translation for dropseq-tools, the cell ID and UMI values were assigned to 12-bp and 8-bp random sequence segments, respectively. For the UMI bequeathing strategy, the source UMI sequences were elongated by A nucleotides to adjust the UMI lengths if necessary. The process configuration files of INTERSTELLAR are available at https://github.com/yachielab/Interstellar/tree/main/config.

### scRNA-seq data analysis

We translated scRNA-seq read datasets of 10X Chromium V3 scRNA-seq, Drop-seq, Quartz-seq2 v3.1 and SPLiT-seq and analyzed them by 10X Cell Ranger (version 3.0.2) and dropseq-tools (version 2.3.0). For comparison, we also analyzed the original read datasets by their proprietary software tools (i.e., 10X Cell Ranger, dropseq-tools, Quartz-seq pipeline, and splitseq-tools (commit c3923ea), respectively). The mouse reference genome GRCm38 was commonly used throughout these analyses. In the analysis of both the original SPLiT-seq read dataset and those translated and analyzed by the other two software tools, cDNA segments mapped to intronic regions were also accounted for to estimate gene expression. Gene expression count matrices obtained from the original software tools were processed to filter out low-quality cells with the following criteria using Seurat version 3.2.0^42^. For those from 10X Cell Ranger, dropseq-tools, and Quartz-seq pipeline, we removed cells whose numbers of detected genes were ≤200 or ≥2500 or whose UMI proportion from mitochondrial genes was ≥20%. For the original gene expression matrix of SPLiT-seq obtained by splitseq-tools, we removed cells whose numbers of detected genes were ≤250 or ≥2500 or UMI proportion from mitochondrial genes was ≥5%, considering the expectation of low mitochondrial reads in the single-nucleus RNA-seq. Finally, cells commonly observed in the gene expression matrices obtained by the original software tool, 10X Cell Ranger, and dropseq-tools were retained, yielding 5,003, 11,334, 1,048, and 185,722 cells for ones sourced from the original 10X Chromium V3, Drop-seq, Quartz-seq2, and SPLiT-seq datasets, respectively. Using Seurat, all of the filtered gene expression matrices were then processed by NormalizeData() with a scale factor of 10000 and ScaleData(), followed by the extraction of the top 5000 highly variable genes by FindVariableFeatures() for principal component analysis (PCA) by RunPCA(). Using the top 20 principal components, we carried out two-dimensional UMAP embedding and kNN clustering of each dataset by RunUMAP(), FindNeighbors(), and FindClusters() with the resolution parameter of 0.6.

### Simulation of a synthetic scRNA-seq read structure

We simulated a sequencing read pool of a highly complex read structure from a total of four 10X Chromium V3 scRNA-seq read datasets (two heart and two neural samples) obtained from the 10X Genomics website. Since these datasets had already been demultiplexed, we provided simulated Illumina i5 and i7 index reads in addition to each paired-end read entry for cell ID, plus UMI and cDNA. The i5 indices were provided to discriminate individual FASTQ datasets, and the i7 indices were provided to indicate the sample type (heart or neural), in which they could serve as parental sequence segments of their associated i5 indices. The sequencing read datasets were pooled and interpreted by INTERSTELLAR as described above for 10X Chromium V3 reads. Next, the sequencing reads were translated into a destination read structure with the value space optimization process, where 15-bp i7 index, 10-bp i5 index, 16-bp cell barcode, and 12-bp UMI segment sequences were translated into arbitrarily designed three 2-bp units, four 3-bp units, five 4-bp units (each unit was limited to a selection from ten 4-bp sequences in an allowlist) and five 2-bp units, respectively. Finally, the simulated reads were translated back to the read structure of 10X Chromium V3. The original and round-trip reads were both analyzed by 10X Cell Ranger. The gene expression matrices were derived by the same criteria described above. The process configuration file of INTERSTELLAR is available at https://github.com/yachielab/Interstellar/tree/main/config/figS2_10X.

### Translation of spatial transcriptomics reads

Using INTERSTELLAR, we analyzed Slide-seq reads and identified their positional barcodes, UMIs, and cDNA fragments. We discarded sequencing reads with any segment whose average Q score was below 20 or minimum per base Q score was below 5 and reads whose positional barcodes were not found in the allowlist with the perfect match. After obtaining a sequence conversion table between positional barcodes of a Slide-seq slide and those of multiple 10X Genomics Visium slide tiles, reads were grouped by destination Visium tile. For each Visium tile group, we translated the reads into the Visium read structure using INTERSTELLAR with the UMI bequeathing strategy, where the segment length is adjusted by adding A nucleotides. The process configuration file of INTERSTELLAR is available at https://github.com/yachielab/Interstellar/tree/main/config/fig5_Slide-seq. The resulting FASTQ files of Visium tiles were independently processed by 10X Genomics Space Ranger (version 1.0.0) with the options “--slide=V19L01-041 --area=C1” using a fake slide image (https://github.com/yachielab/Interstellar/blob/main/utils/fake_spaceranger_box.jpeg) such that whole Visium spots were recognized to be covered by a tissue sample and processed. For each Slide-seq tissue sample, the Space Ranger results of multiple tiles were merged and analyzed by Seurat version 3.2.0 to obtain a single gene expression count matrix of spatial positions, with the same protocol applied for the scRNA-seq data analyses above, except that the top 3000 highly variable genes were used for PCA and the top 30 principal components were used for UMAP embedding and kNN clustering.

### Translation and analysis of sci-Space data

Using INTERSTELLAR, we analyzed sci-Space reads of slide IDs 7–14 listed in Supplementary Table 1, identified their cell ID and UMI segments, and translated the reads into a single pair of FASTQ files for 10X Cell Ranger with the UMI bequeathing strategy. The process configuration file is available at https://github.com/yachielab/Interstellar/blob/main/config/fig5_sci-Space/sciSpace.conf. After analyzing the translated reads by Cell Ranger, we processed the expression matrices obtained from the original pipeline (https://ftp.ncbi.nlm.nih.gov/geo/series/GSE166nnn/GSE166692/suppl/GSE166692_sciSpace_count_matrix.mtx.gz) and Cell Ranger with the same criteria used for the scRNA-seq analysis above. From each dataset, we independently performed kNN clustering and obtained cell state labels. Cell state occupancies in each spot were plotted as pie charts using an R package scatterpie (https://github.com/GuangchuangYu/scatterpie) based on the coordinate information of cells from the original study (https://ftp.ncbi.nlm.nih.gov/geo/series/GSE166nnn/GSE166692/suppl/GSE166692_sciSpace_cell_metadata.tsv.gz).

### Statistical tests

The Euclidean distance correlations in high-dimensional data space between the original and translated results were all measured by Pearson’s correlation. The statistical tests to compare the rank difference distributions to random expectations were performed by the two-sided Wilcoxon rank-sum test.

### Code availability

INTERSTELLAR is available at https://github.com/yachielab/INTERSTELLAR. All the codes used in this study are available at https://github.com/yachielab/Interstellar/tree/main/utils. Test codes are executable at https://colab.research.google.com/drive/1nuqPK_zQSXFXHu-9gZR5w9EfsQhH6Itl?usp=sharing.

### Data availability

The RCP-PCR data generated in this study have been submitted to the NCBI BioProject database (https://www.ncbi.nlm.nih.gov/bioproject/) with the accession number PRJNA767068.

## Supplementary Materials for

**Extended Data Figure 1.**
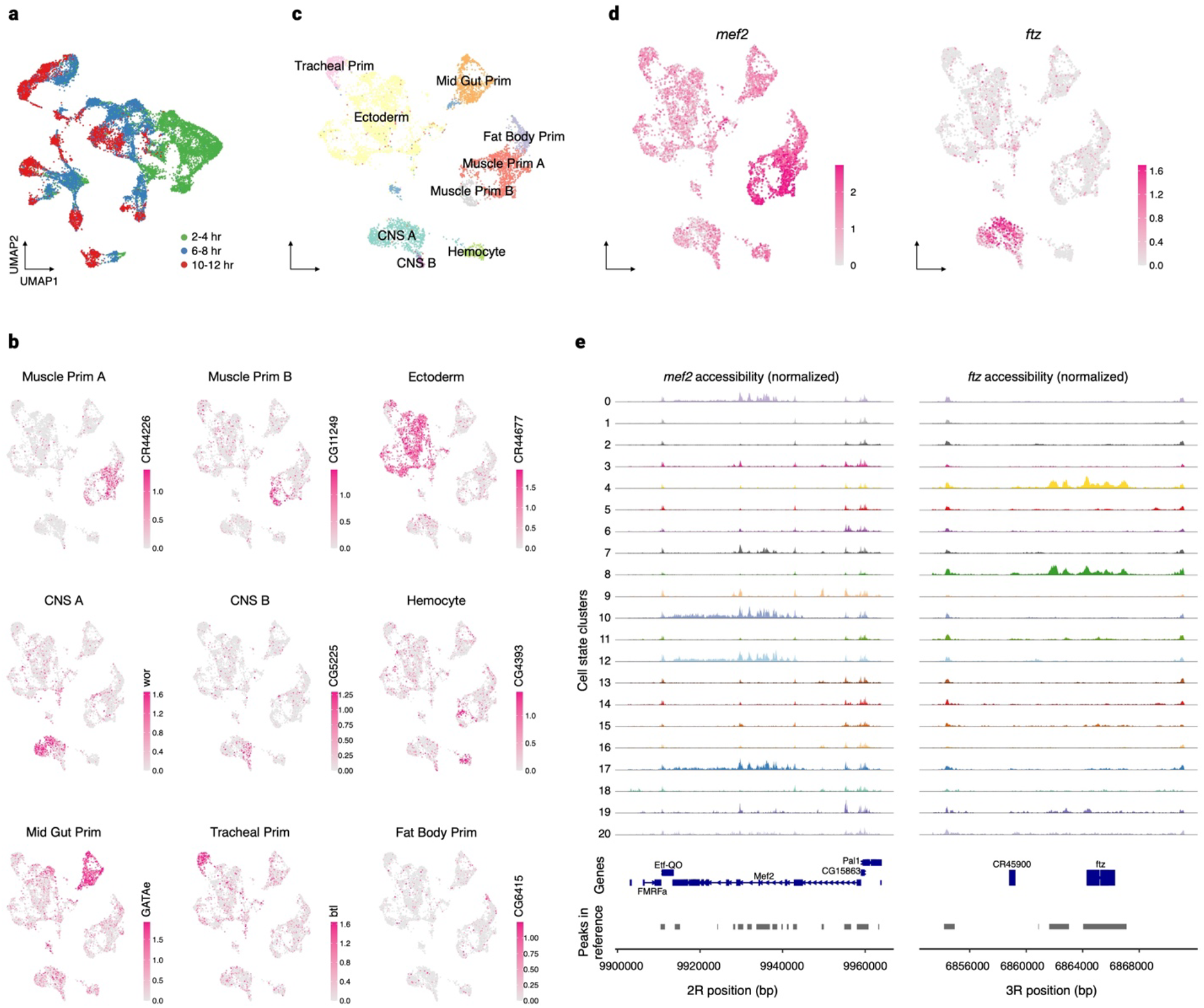
Analysis of sci-ATAC-seq reads translated for 10X Cell Ranger ATAC. **a**, Two-dimensional UMAP embedding of the sci-ATAC-seq datasets processed by its original pipeline for Drosophila embryo 2 to 4, 6 to 8, and 10 to 12 hours after egg laying. **b-e**, Analyses of the same dataset translated by INTERSTELLAR and analyzed by 10X Cell Ranger ATAC. **b**, Single-cell chromatin accessibilities of marker genes. **c**, Cell state annotations. **d**, Single-cell chromatin accessibilities of *mef2* and *ftz*. **e**, Chromatin accessibilities of *mef2* and *ftz*-encoding regions in different cell state clusters represented in Fig. 3b.

**Extended Data Figure 2.**
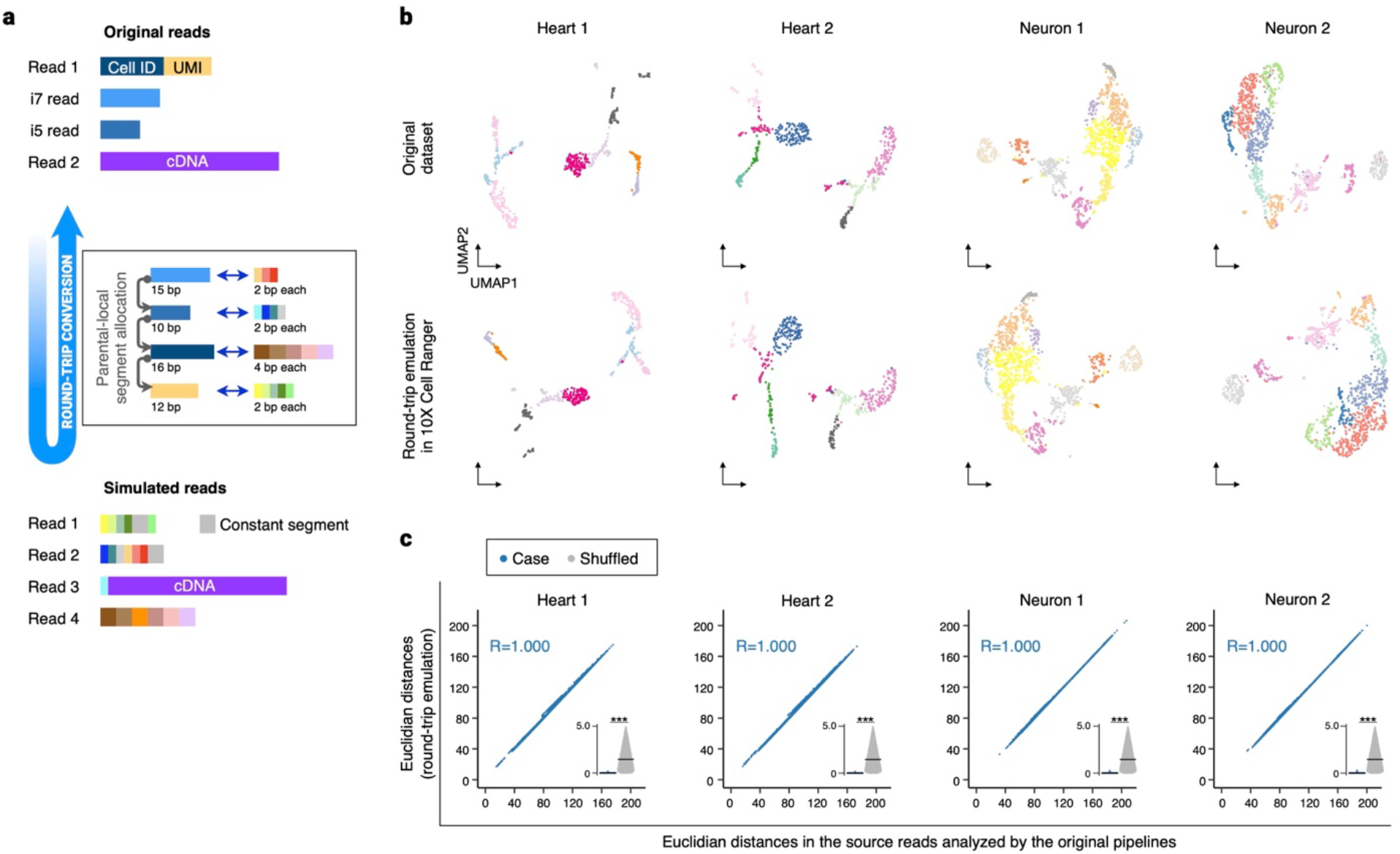
A round-trip conversion between two read structures. **a**, Four 10X Chromium V3 scRNA-seq libraries each with a unique library index pair were combined and transformed into a simulated pooled library of a complex read structure using INTERSTELLAR. The simulated library was then translated back into a pooled library of the Chromium V3 read structure. **b**, Two-dimensional UMAP embeddings of four original scRNA-seq datasets (Heart 1K Lane 1, Heart 1K Lane 2, Neuron 1K Lane 1, and Neuron 1K Lane 2) and those by the round-trip conversion. Cell cluster annotations of single cells obtained by the original pipelines were applied to the round-trip conversion results. **c**, Correlation in Euclidean distance of two cells in high-dimensional transcriptome space between the original datasets and those produced by the round-trip conversion. For each dataset, Euclidean distances in the gene expression count matrix were measured for 50,000 randomly sampled cell pairs. The inset sina plots represent rank difference distribution in the Euclidean distance of the same cell pairs before and after translation. The crossbar represents the median.

